# Copper nanoparticles as modulator of active bacterial population and their physiology in ecosphere

**DOI:** 10.1101/2021.11.30.470660

**Authors:** Ali Alboloushi, Absar Alum, Morteza Abbaszadegan

## Abstract

The widespread use of nanoparticles (NPs) in industrial and consumer products has resulted in their emergence as significant environmental contaminants that can potentially modulate the role of bacteria in environment. This study examines the impact of different sizes of Copper nanoparticles (CuNPs) on the population and physiology of environmentally relevant gram positive (*Bacillus)* and gam negative (*Alcaligenes*, and *Pseudomonas*) bacteria. In general, exposure to CuNPs resulted in 4 to >6 log inactivation in bacterial population. More specifically, after 2hr exposure of *Alcaligenes* and *Pseudomonas* to 50 CuNPs, 5.75 and 6.64 log reduction noted, respectively; and their exposure to 100 CuNPs resulted in 5.97 and 6.58 log reduction, respectively. A similar exposure of laboratory and environmental isolates of *Bacillus* to 50 and 100 CuNPs resulted in 4.84, 4.16 and 4.35, 3.61 log reduction, correspondingly. The exposure induced elicitation of different toxicity pathways in the test bacteria. Bacterial exposure to 50 CuNPs resulted in elevated levels of LDH in *Pseudomonas*, in contrast these levels decreased in *Alcaligenes* and *Bacillus*. Our toxicity studies showed that exposure to CuNP can have various levels of metabolic and cellular modulation in bacterial species, suggesting that the presence of CuNPs in environment can potentially impact the pollutants-attenuation-role of bacteria in environments such as wastewater biological treatment processes.

## Introduction

Widespread use of nanoparticles (NPs) in agricultural, industrial and consumer products results in the release of huge quantities of NPs in environment. The NPs species have been recognized as significant contaminants in wastewater and sludge (EPA 2003; Roco 2007; Westerhoff et al., 2009). They are known to impact biological activities at molecular, cellular, organism/system??, and ecological levels (Aruoja, et al., 2015; Bhuvaneshwari, 2016; Joonas et al., 2019; Ren et al., 2009). Unabated release of heavy metal-containing nanoparticles (mNPs) into the wastewater can have detrimental consequences on the normal bio-ecological processes. Increased prevalence of mNPs can potentially alter a range of ecologically beneficial roles played by bacteria, including the normal function of wastewater treatment processes, natural attenuation of pollutants and mineral cycling. The functional role of mNPs in the ecological processes have been broadly studied under laboratory and small-scale field studies; there is a need for investigating these impacts under well-defined experimental conditions (Alum et al., 2018).

Copper nanoparticles (CuNPs) are widely used in industrial applications such as semiconductors, heat transfer fluids in machine tools, metal catalysts, and even in the area of public health as biocidal preparations (Aruoja et al., 2009; Kim et al., 2011). In addition, a large amount of CuNPs is continuously released from the automobile brake systems (Denier et al., 2007; Grigratos and Matini 2015). These facts make CuNPs a good candidate for studying their potential adverse effects on bio-geo-ecology of surface and waste waters (Brown et al., 1995; Gilbertson et al., 2016).

The objective of this study was to investigate the adverse impacts of CuNPs on the laboratory isolates of *Bacillus, Alcaligenes*, and *Pseudomonas* and field isolates of *Bacillus*. The study also measured the role of various environmental conditions, such as pH, temperature, and light exposure on the mNPs role in environment.

## Materials and Methods

### Bacterial Strains

Pure cultures of *Bacillus subtilis* (ATCC 23059), *Alcaligenes faecalis* (ATCC 8750), and *Pseudomonas aeruginosa* (ATCC 10145) were obtained from the American Type Culture Collection (ATCC, Rockville, MD). A *Bacillus* isolate from a local wastewater lagoon (Maricopa, AZ) was included in the study as a representative of environmental isolates for comparison purposes. The identity of environmental isolate was confirmed by molecular analyses such as polymerase chain reaction (PCR) and sequencing (data not shown).

### Culture Preparation

Working cultures of the selected bacteria were prepared by inoculating 0.1 ml of the over-night culture in 9.0 ml of the nutrient broth and incubated in a shaker-incubator (New Brunswick Scientific C24, Edison, NJ) (150 RPM @ 37°C). Cultures were grown to an optical density of 0.8 to 1.0 at 600 nm, measured using a spectrophotometer (Hach DR/4000U, Loveland, CO). Bacterial cells were harvested by centrifugation at 1500xg for 10 min; supernatants were discarded, and pellets were re-suspended in phosphate buffer (0.5M PBS).

### Characteristics of Copper Nanoparticles

The CuNPs included in this study were spherical in shape with a diameter of 50 nanometer (50 CuNP) (Cat# 684007) and 100 nanometers (100 CuNP) (Cat# 634220) (Sigma Aldrich, Saint Louis, MO). Both types of CuNPs species have similar physicochemical characteristics including density (8.94 g/mL); resistivity (1.673 μΩ-cm at 20°C); surface area (5-10 m^2^/g) and surface charge (+33mV). Identical physicochemical properties of both test CuNPs permitted to delineate the impact of particle size on their biological impact. Stocks of test CuNPs were kept at room temperature and suspended in PBS as needed. The distribution of CuNPs in reactor solution was determined by Dynamic Light Scattering technique and polydispersity index of 0.225 and 0.057 was recorded for sonicated stock and reactor sample.

### Bacterial Exposure to CuNPs

Exposure experiments were conducted in an axially rotating mixer (Rotamix, ATR Inc. Laurel, MD) operated at 40 RPM to ensure constant and even dispersal of CuNPs in the reaction volume throughout the experiment. In the exposure studies, use of the axially rotating mixer is critical as CuNPs tend to settle down in the peripheral zone of a stir mixer.

Each type of NPs (50 CuNPs and 100 CuNPs) was added to the individual reactor at a final concentration of 6mM (0.0036 mg per10 mL).

Total number of 50 CuNPs in each reactor = (total mass) × (density)^-1^ × (volume)^-1^ = (0.0036 mg) × (1×10^21^ nm^3^) × (8.94)^-1^ × (1,000 mg)^-1^ × (6.5×10^4^ nm^3^)^-1^ = 6.2×10^9^ 50 CuNPs

Total number of 100 CuNPs in each reactor = (total mass) × (density)^-1^ × (volume)^-1^ = (0.0036 mg) × (1×10^21^ nm^3^) × (8.94)^-1^ × (1,000 mg)^-1^ × (5.2×10^5^ nm^3^)^-1^ = 7.7×10^8^ 100 CuNPs

Exposure experiments were conducted in individual reactors containing 50 or 100 nm CuNPs spiked with washed bacteria at a concentration of 1×10^9^ CFU/mL. The control reactor was operated under similar experimental conditions with no CuNPs added. Samples were collected at 20, 40, 60, 90, and 120 min contact time from each reactor and analyzed in duplicate assays for viable bacterial counts using membrane filtration.

### Membrane Filtration for Bacteria

Samples were analyzed for target bacteria by filtering samples through a 47 mm cellulose acetate membrane with 0.47 µm pore size. Each membrane was placed on appropriate selective agar media plate and incubated at 37°C for 24-48 hours. After the incubation, colonies were counted and recorded as viable bacterial counts for each exposure time.

### CuNP Toxicity Pathways

The toxicity of CuNPs was studied by examining their; 1) interaction with the ligands on bacterial cell surface; 2) impairment cellular defense mechanisms; 3) interference with the cellular energy pathways. Washed bacterial cells were exposed to CuNPs for a specified time and exposed cells were analyzed for the metabolic activities including lactic dehydrogenase, glutathione reductase enzyme and NAD(P)H-dependent oxidoreductase.

### Lactic Dehydrogenase Assay

After exposure to CuNPs, damage to bacterial cell membrane was determined using the Lactic Dehydrogenase based Cellular In Vitro Toxicology Assay Kit, (Sigma-Aldrich, Saint Louis, MO). The assay was performed according to the manufacturer’s instructions with slight optimization for detecting the total LDH in bacteria cells and samples suspension.

### Glutathione Reductase Assay

The glutathione reductase (GR) enzyme protects cells against radicals. The level of GR in CuNPs exposed cells were determined using the Glutathione Assay Kit, (Cayman Chemical Company, Ann Arbor, MI). The analyses were performed according to the manufacturer’s instructions. The absorbance readings for GR activity were normalized using the equation provided in the GR assay protocol:

The actual extinction coefficient for NADPH at 340 nm adjusted for the path length of the solution in the cuvette = 0.00622 μM^-1^cm^-1^x1cm = 0.00622

### MTT Assay

The integrity of energy cycle in the CuNPs exposed bacterial cells was determined by quantify the activity of NAD(P)H-dependent oxidoreductase. The assay was performed using the MTT Cell Proliferation Assay Kit obtained from American Type Culture Collection (ATCC) (Manassas, VA). The kit was used according to the manufacturer’s instructions.

### Scanning Electron Microscope

The physical impact of 50 & 100 nm CuNPs on the *Bacillus* cells was visualized using scanning electron microscope (SEM) (Focused Ion Beam – Nova 200 – NanoLab - FEI) (Hillsboro, OR). The bacterial cells exposed to 50 & 100 nm CuNPs were deposited on aluminum stud and coated with gold using sputter coater (Denton Vacuum) (Moorestown, NJ). The samples were observed under the scanning electron microscope to locate CuNPs inside *Bacillus* cells. As a control, non-treated *Bacillus* cells were identically processed and visualized. The treated and control samples were observed under SEM with XLD scan capabilities that allowed to detect the CuNPs inside the bacterial cell using Cu specific peaks in XLD scans.

### Release of Cu^2+^ from CuNPs

The kinetics of the Cu^2+^ release from 50 & 100 nm CuNPs in aqueous solution (0.5M PBS buffer) was studied using Cupric Ion Selective Electrode (Cu^2+^-ISE) (Cole-Parmer, Vernon Hills, Illinois). The release of Cu^2+^ were studied under different conditions including light exposure, temperature, contact time, and medium acidity. The Cupric Ion Selective Electrode was used according to the manufacturer’s instructions. Cu^2+^ complex formation was studied in 0.5M PBS and nano-pure water and calibration curves for both solutions were generated.

## Results and Discussion

### Bacterial Inactivation by CuNPs

The impact of 50 & 100 nm CuNPs on environmentally important gram-negative and gram-positive bacterial isolates was investigated. After short exposures to 50 and 100 CuNPs, *Bacillus, Alcaligenes*, and *Pseudomonas* showed differences in inactivation kinetics; however, after 120 min exposure, the cumulative bactericidal effect of both type of CuNPs appears to reach the same rate (Figure 1). The highest bactericidal effect of 50 and 100 CuNPs against laboratory isolates of *Alcaligenes*, and *Pseudomonas* was noted at 90 min. Whereas, for *Bacillus* the bactericidal effect peaked at 60 and 40 min for 50 and 100 CuNPs, respectively (Table 1).

**Figure. 1.**
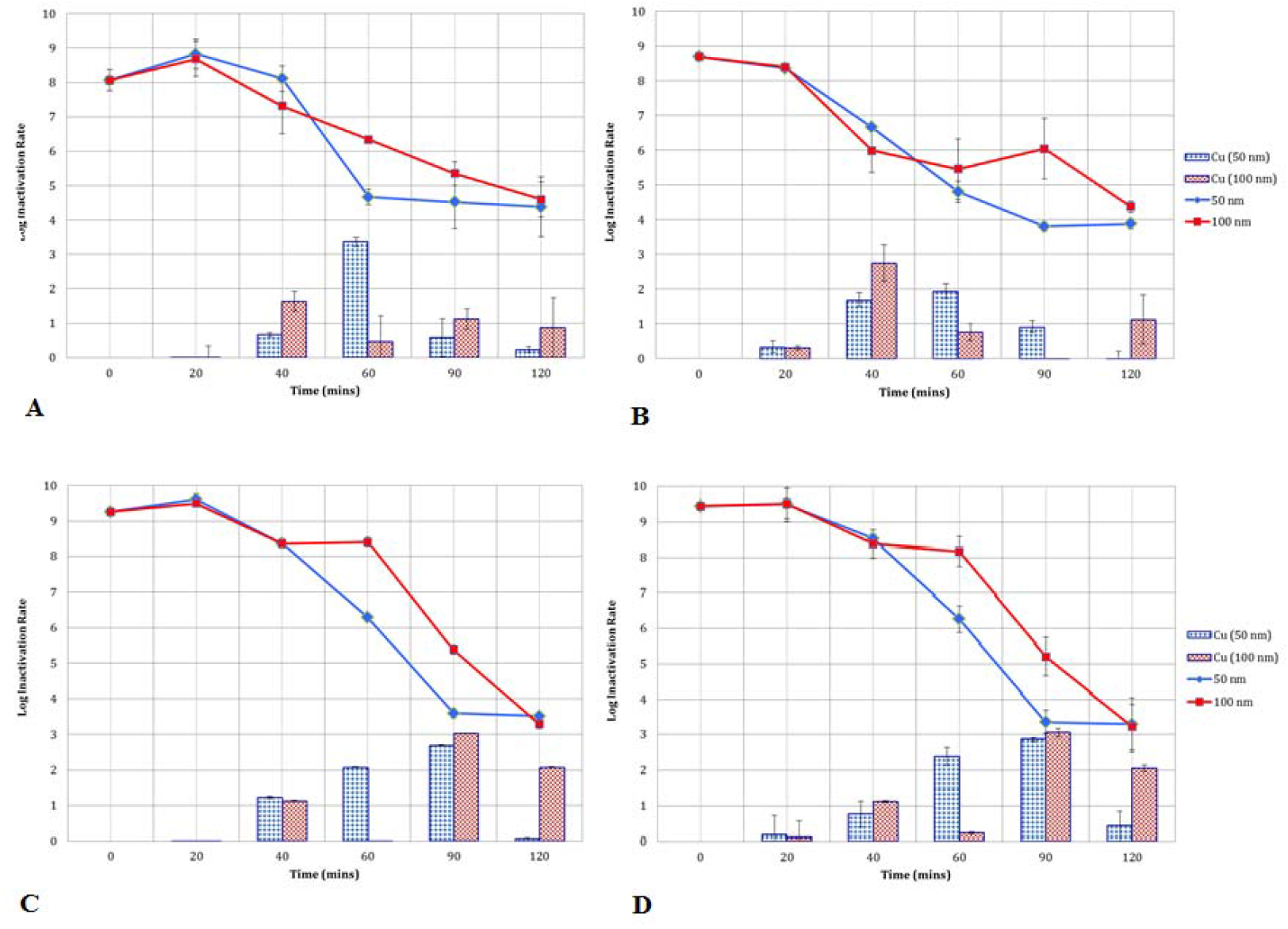
Bactericidal effects of 50 and 100 CuNPs on: A) *Bacillus* the laboratory isolate, B) *Bacillus* isolate from wastewater, C) *Alcaligenes* laboratory isolate, D) *Pseudomonas* laboratory isolate. ***Note:*** The line graphs represent accumulative inactivation and the bar graphs represent inactivation between time intervals.

**Table 1.** Inactivation of laboratory and environmental isolates of G+ and G-bacteria using 50 and 100 CuNPs

| bacteria using 50 and 100 CuNPs |  |  |  |  |
| --- | --- | --- | --- | --- |
| Bacterial Species | Peak Inactivation (min) |  | Total Reduction Log<sub>10</sub> |  |
|  | 50 CuNPs | 100 CuNPs | 50 CuNPs | 100 CuNPs |
| *Bacillus* (Laboratory Isolate) | 60 | 40 | 4.16 | 3.61 |
| *Bacillus* (Wastewater Isolate) | 60 | 40 | 4.84 | 4.35 |
| *Alcaligenes* (Laboratory Isolate) | 90 | 90 | 5.75 | 5.97 |
| *Pseudomonas* (Laboratory Isolate) | 90 | 90 | 6.64 | 6.58 |

### Biocidal efficacy of CuNPs in Wastewater

The bactericidal efficacy of 50 and 100 CuNPs were investigated in primary and secondary treated wastewater samples. In primary treated wastewater, after 2hr exposure to 50 or 100 nm CuNPs, the concentration of bacteria decreased from 6.3 to 2.7 & 6.3 to 3.1 logs, respectively (Figure 2). Whereas, in secondary treated wastewater, the concentration of bacteria decreased from 3.9 to 2.1 & 3.9 to 2.3 logs, respectively (Figure 2). In general, 50 CuNPs showed comparatively greater bactericidal effects than the100 CuNPs. Besides NPs size, the exposure conditions also appear to influence the antibacterial efficacy of CuNPs. Discontinuation of mixing resulted in increased bacterial concentrations in both primary and secondary treated wastewater samples. The high organic matter contents in these water samples supported the bacterial regrowth. However, resumption of mixing after 72 hr, again resulted in declined concentration of bacteria in both samples. The results highlight the role of the appropriate mixing conditions that ensures constant contact between the bacterial cells and the biocidal CuNPs.

**Figure 2.**
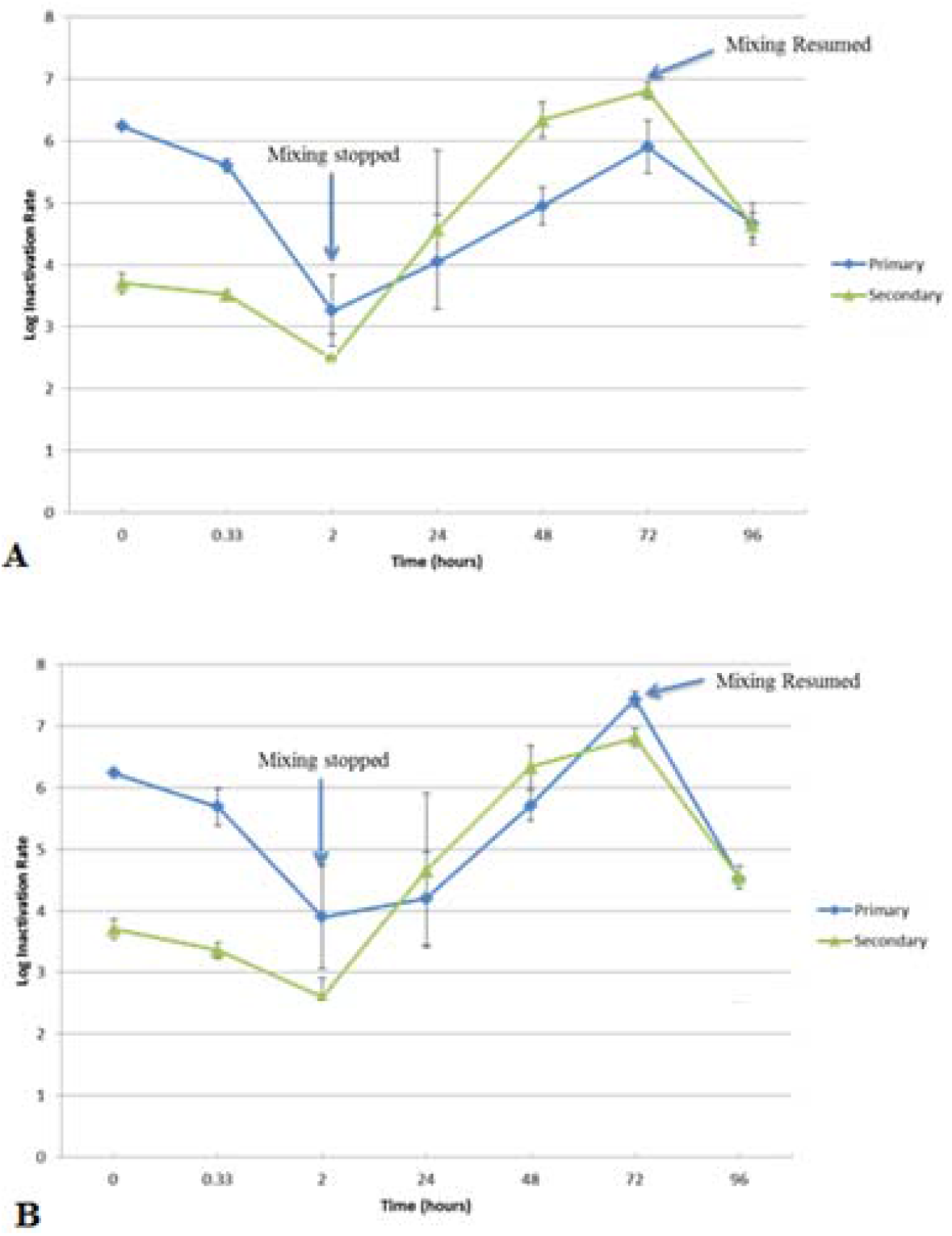
Effects of mixing conditions on the antibacterial efficacy of CuNPs in the primary and secondary treated wastewater samples, A) 50 CuNPs, B) 100 CuNPs.

Furthermore, the antibacterial impact of 50 CuNPs on *E. coli* in the sterilized primary and secondary treated wastewater samples was investigated. After 2 hr mixing with 50 CuNPs, *E. coli* concentration in the sterilized primary and the sterilized secondary treated wastewater decreased by 4.5 and 3.4 log, respectively (Figure 3). However, under both conditions, the concentration of *E. coli* increased after mixing was stopped for 24 hrs, and it continued to increase until mixing was restarted. This increase in bacterial numbers is thought to be due to the high level of organic matter in these samples which might have supported bacterial regrowth. Termination of mixing resulted in a rapid deposition of CuNPs at the bottom of the reactor while bacterial cells remained suspended in solution. The fact that antibacterial effect against *E. coli* is observed only under appropriate mixing condition, suggesting that CuNPs physical proximity to bacterial cells is critical for exerting bactericidal effect. Presence of CuNPs sediments at the bottom of the reactor did not have any bactericidal impact on planktonic bacterial population suspended in solution. The results clearly highlight the importance of appropriate mixing conditions under laboratory conditions and turbulence in aquatic system for continued biocidal impact of CuNPs in ecosystem.

**Figure 3.**
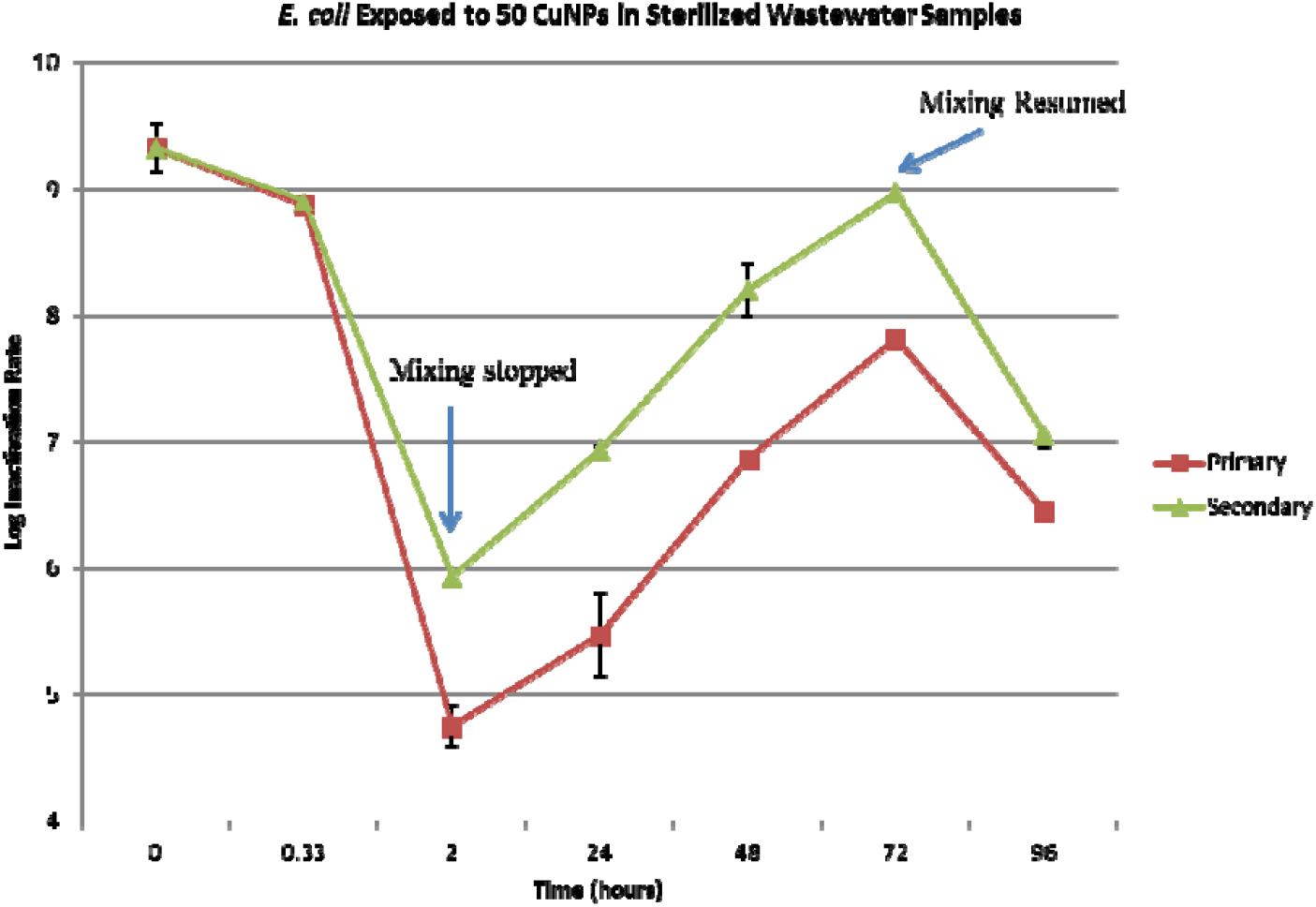
Impact of mixing conditions on the biocidal efficacy of 50 CuNPs against *E. coli* in the sterilized primary and secondary treated wastewater samples.

### Toxicity Pathways

#### LDH Assay

CuNPs toxicity against gram+ and Gram-bacteria was studied using LDH assay. Exposure to 50 CuNPs resulted in decreased LDH levels in all the bacterial isolates investigated in this study except for *Pseudomonas*, (Table 2). The total LDH activities inversely correlated to the total viable cells remaining in the sample at any specified time. In the laboratory isolate of *Bacillus*, slightly lower LDH activities were noted compared to other laboratory bacterial isolates. This could be due to the difference in the stress response pathway used by bacterial strains to handle elevated levels of copper in cell. For example, copper stress results in the expression of copper-binding proteins (CuBPs) on the surface of some bacterial species such as *Pseudomonas* and *Vibrio;* however, no such CuBPs are expressed on the surface of *Bacillus* (Rensing and Gross, 2003). Therefore, further molecular work is required to determine exact nature of such responses by different bacterial strains.

**Table 2:** Elicitation of LDH activities in bacterial strains exposed to 50 and 100 CuNPs for 120 min.

|  |  |  |
| --- | --- | --- |
| Strain | 50 CuNPs | 100 CuNPs |
| *Bacillus* (Lab. Isolate) | ↓ | ↓ |
| *Bacillus* (Field Isolate) | ↓ | ↓ |
| *Alcaligenes* (Lab. Isolate) | ↓ | ↓ |
| *Pseudomonas* (Lab. Isolate) | ↔ | ↓ |

Note: ↓= decrease. ↔= No change

#### GR Assay

Impact of CuNPs exposure on glutathione reductase levels in the bacterial cells was studied. Exposure to 100 CuNPs resulted in elevated GR activities in field and laboratory isolates of *Bacillus*; however, exposure to 50 CuNPs caused such effect only in field isolate of *Bacillus* and not the laboratory isolates (Figure 4). Similarly, exposure to 100 CuNPs provoked elevated GR activity in *Alcaligenes* cells; however, exposure to 50 CuNPs did not produce such response. In general, the GR activities declined over exposure time. In case of *Pseudomonas*, GR activities remained unchanged after exposure to 50 and 100 CuNPs. The variations in GR induction levels among different species could be caused by many factors. For example, part of the glutathione may be derived from the culture medium consisting yeast extracts that are ordinarily rich in glutathione. Even though samples were washed, it is possible some glutathione from medium may still be attached to bacterial cells and interfered with the GR determinations. In addition, GR activities are species-specific, a wide range of bacteria lack glutathione while others may have a very small concentrations (Fahey et al., 1978).

**Figure 4.**
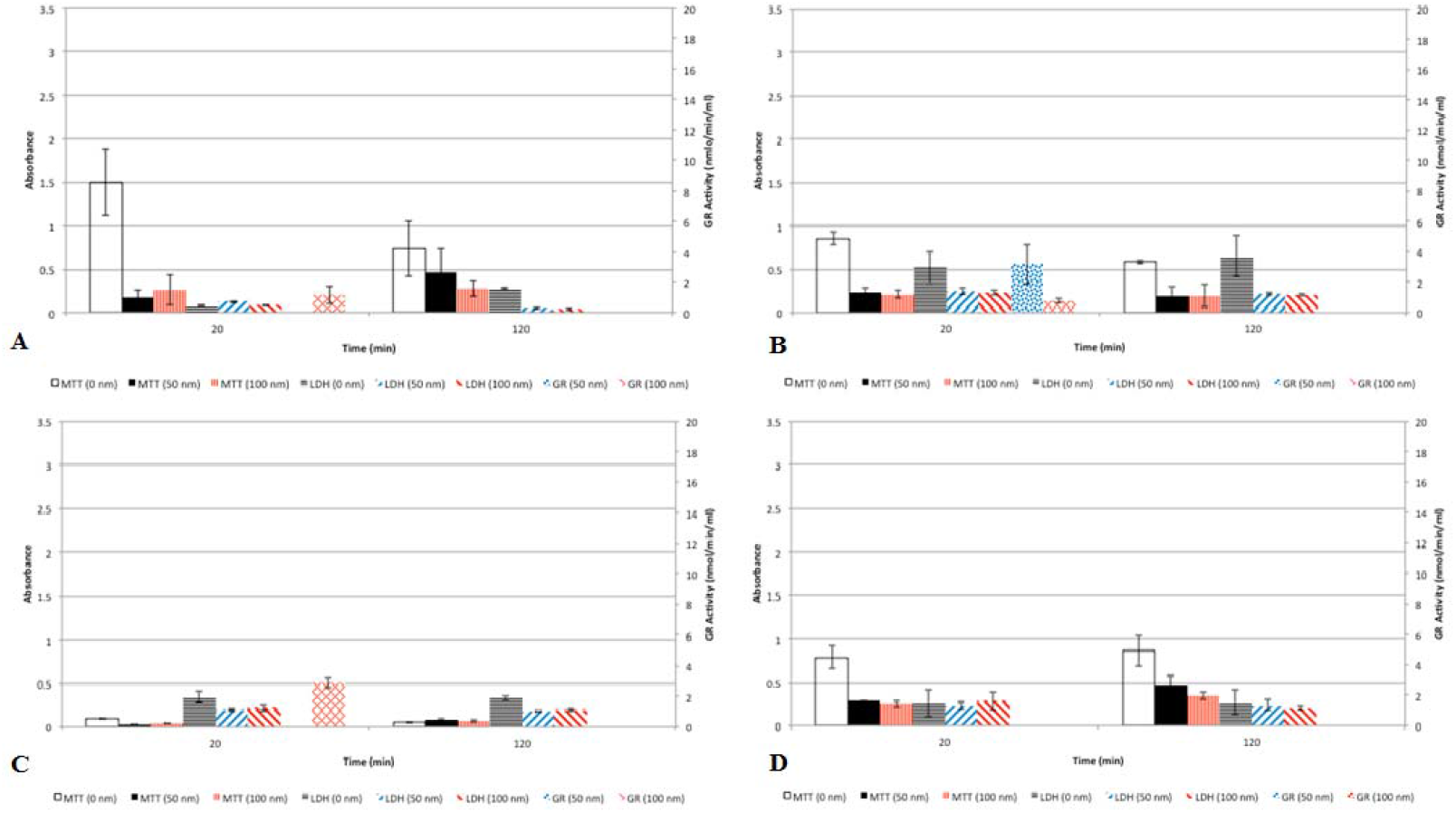
Metabolomic profile of bacteria after 120 min exposure to CuNPs, A) *Bacillus* laboratory isolate, B) *Bacillus* environmental isolate, C) *Alcaligenes* laboratory isolate, D) *Pseudomonas* laboratory isolate.

#### MTT Assay

The impact of CuNPs exposure on bacterial energy metabolic pathways was studied using MTT as endpoint. In general, with increasing exposure time, the MTT activity increased in laboratory isolates of *Bacillus, Alcaligenes, and Pseudomonas* and decreased in field isolate of *Bacillus* (Figure 4). The levels of MTT activity in *Alcaligenes* cells were relatively low compared to the other bacterial strains included in this study. These results indicate that CuNPs may impact at different levels of energy pathway in different bacterial isolates.

#### Release of Cu^2+^ from CuNPs

The release of Cu^2+^from CuNPs in aqueous solution (0.5M PBS buffer) were examined and no detectable concentrations of Cu^2+^ were recorded under the conditions tested in this study (Table 3). Even extended exposure up to 200 hours did not result in release of any detectableCu^2+^; however, a light-blue precipitate was formed at the bottom of the solution (Data not shown). The blue precipitate is copper hydroxide (Cu(OH)_2_,) which is formed by the reaction of Cu^2+^ ions with OH^-^ ions in water. This is an indication of Cu^2+^ ions release from CuNPs in aqueous solution; however, it is assumed that they are released at a slow rate and instantly react with OH^-^ ions in water, minimizing their possible role in bacterial inactivation.

**Table 3:**
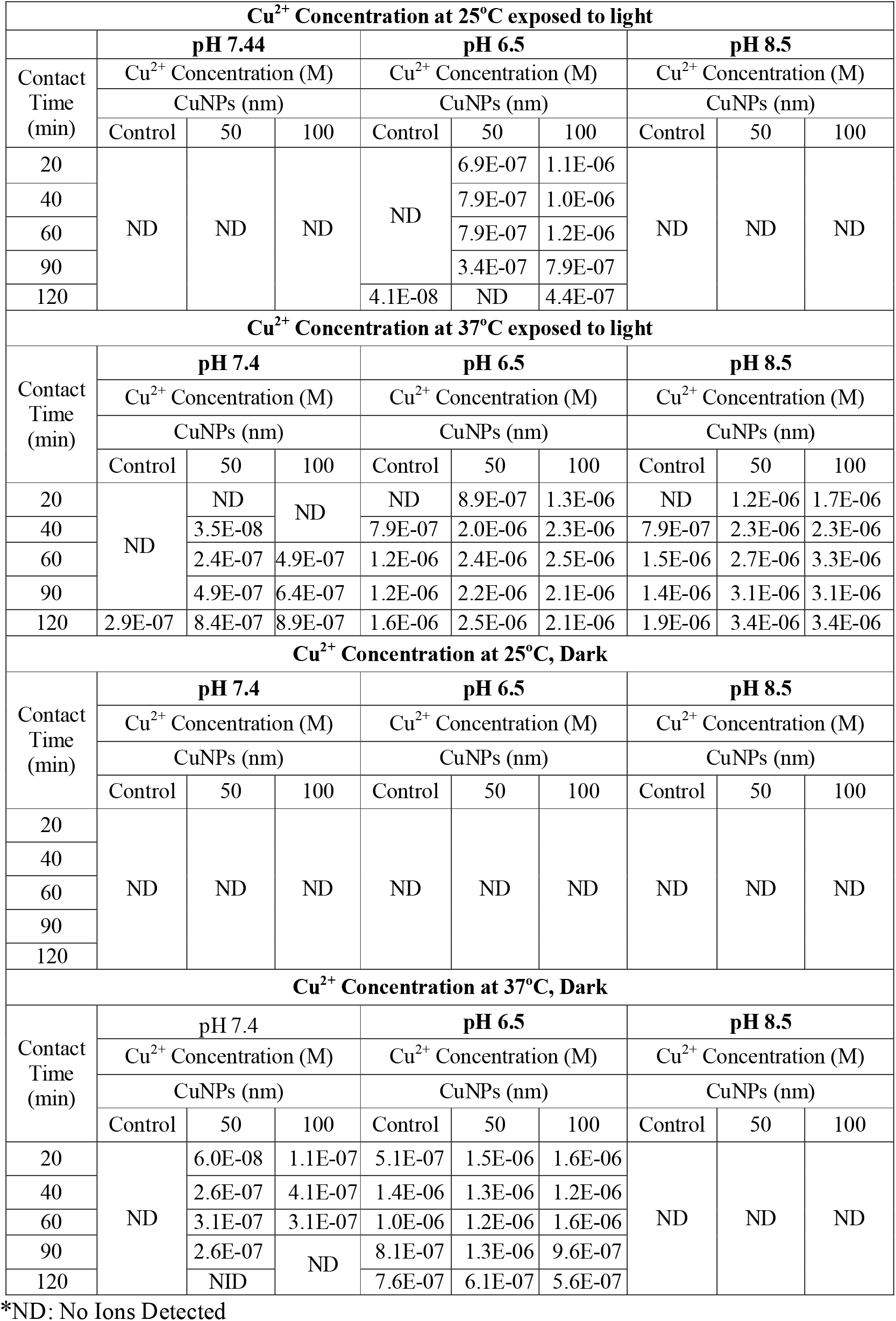
Kinetics of Cu^2+^ release from 50 & 100 nm CuNPs in 0.5M PBS

## Conclusions

CuNPs play a significant role in many new technologies and their use has expanded rapidly over the last recent decades. Most studies on environmental impact of CuNPs have been focused on laboratory strains of bacteria. This study focuses on comparison between laboratory and wastewater isolates of gram-positive and gram-negative bacteria. Subtle differences in the response of laboratory and environmental bacterial isolates and inter-species variations in response to CuNPs were recorded. The study identifies differences in the toxicity elicitation pathway of CuNPs in different bacterial species. Additional studies are recommended for further understanding of these pathways by gene silencing studies, which were beyond the scope of this work.

